# Extreme Value Theory for Modeling Category Decision Boundaries in Visual Recognition

**DOI:** 10.1101/2025.09.08.673121

**Authors:** Jin Huang, Deeksha Arun, Terrance Boult, Walter Scheirer

**Affiliations:** Department of Computer Science and Engineering, University of Notre Dame; Department of Computer Science, University of Colorado, Colorado Springs

## Abstract

Several possibilities exist for modeling decision boundaries in category learning, with varying degrees of human fidelity. This paper finds evidence for preferentially focusing representational resources on the extremes of the distribution of visual inputs in a generative model as an alternative to the central tendency models that are commonly used for prototypes and exemplars. The notion of treating extrema near a decision boundary as features in visual recognition is not new, but a comprehensive statistical framework of recognition based on extrema has yet to emerge for category learning. Here we suggest that the statistical Extreme Value Theory [Coles et al., 2001] provides such a framework. In Experiment 1, line segment stimuli that vary in a single dimension of length [Hsu and Griffiths, 2010] are used to assess how human subjects and statistical models assign category membership to a gap region between two categories shown as reference stimuli. A Weibull fit better predicts an observed human shift when moving from uniform to enriched or long tails as reference stimuli. In Experiment 2, more complex 2D rendered face sequences drawn from morphspaces [Folstein et al., 2012] are used as stimuli. Again, the Weibull fit better predicts an observed human shift when reference stimuli are sampled differently. An extrema-based model lends new insight into how discriminative information may be encoded in the brain with implications for the understanding of how decision making works in category learning.

## Introduction

Human category learning allows for boundaries to be learned between different categories of objects and activities that are observed in an environment such that a subject can reliably assign a category label to something that is known. With decades of work in prototype and exemplar learning for category representation [Ashby and Maddox, 2005, Richler and Palmeri, 2014, Collins and Behrmann, 2020], and discriminative and generative strategies for human decision making [Hsu and Griffiths, 2010, Murphy, 2012, Samadani et al., 2014, Sun and Orchard, 2020, Peters et al., 2024], an important open question remains: how are decision boundaries between categories defined? The answer to this question relies on both the data sampled for representation as well as the mechanism for decision making. From early representations in the brain to more abstract forms of categorization, statistical models based on the Gaussian distribution, central tendency, or expected values are a common and seemingly natural choice for describing sensory data. Prototype and exemplar learning have followed this strategy by emphasizing what is assumed to be most representative of a category. Indeed, when one examines the implementations of classification models used for decision making in category learning, they frequently make use of a Gaussian distribution with a mean and variance [Ashby and Alfonso-Reese, 1995, Rosseel, 2002, Griffiths et al., 2007, Griffiths et al., 2011, Tong et al., 2019, Sanborn et al., 2025] to accomplish this. But is there a strong experimental basis for this central tendency assumption? Findings from neuroscience [Freiwald et al., 2009, Groen et al., 2012a, Leopold et al., 2006b, Tsao and Livingstone, 2008], psychology [Leopold et al., 2001b, Scholte et al., 2009, Tanaka et al., 2012], and computer vision [Scheirer et al., 2014, Scheirer, 2017, Zhang and Patel, 2016, Cruz et al., 2025] suggest an alterative strategy where representational resources are focused on the *extremes* of the distribution, which include the samples closest to any decision boundaries.

In this work, we investigate models based on the statistical Extreme Value Theory (EVT) and show that EVT-based models can more accurately describe input data for category learning tasks. Compared to models based on central tendency that focus on the central portion of the distribution, models based on EVT analyze the distribution’s extreme values in the tail regions to establish decision boundaries. The extreme value theorem [Gumbel, 1954, Coles et al., 2001] is the mathematical foundation behind this idea, which limits the functional forms that must be considered and defines the distribution that data near the decision boundaries must follow. Fig. 1 shows the differences between different strategies for defining decision boundaries for category learning. Compared to a binary discriminative model or a per class generative Gaussian model with Bayesian decision making, a generative EVT model supports multiple classes, better modeling accuracy near decision boundaries, the ability to model asymmetric decision boundaries, and the ability to more accurately predict the probability of a rare sample that belongs to a learned category. The latter point is a significant shortcoming of Gaussian models, which tend to underestimate probabilities in such circumstances. The ability of an EVT model to adapt itself to data around decision boundaries allows for decision boundaries to take on different transitions. For instance, in Fig. 1, tail estimate 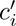, which has three closely spaced points near the decision boundary, has a very sharp transition, while tail estimate 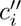 has a slower transition because there is greater spread at the boundary. The research question this work addresses is whether there is any behavioral basis for such an EVT model.

**Figure 1:**
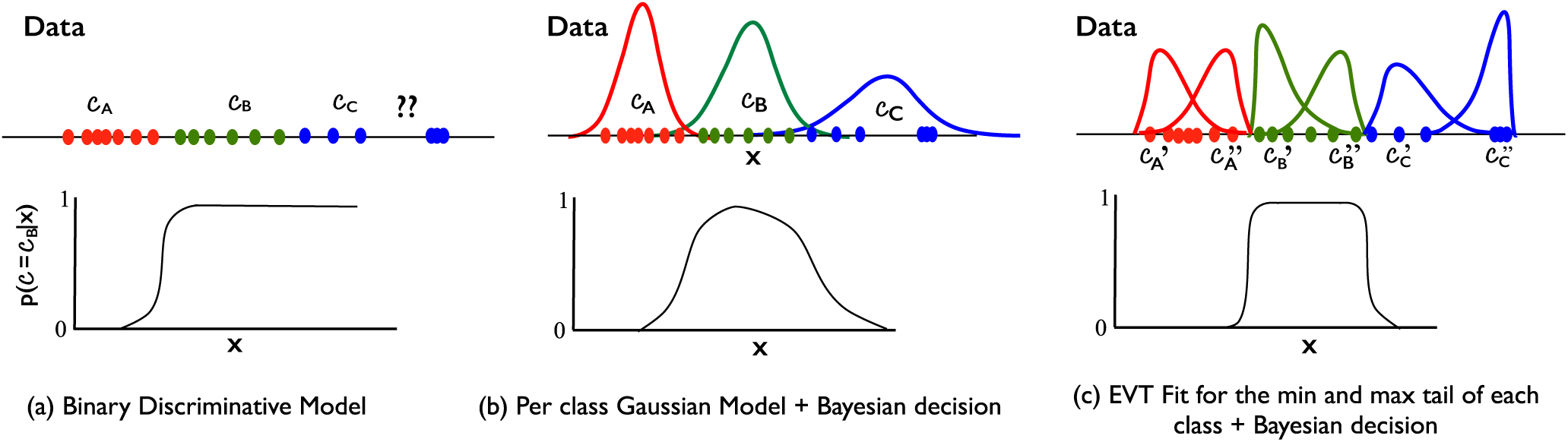
How is an extreme value theory model for category learning different from a central tendency model? The first panel (a) shows data from two known categories, and a hypothetical model that could be learned from that data to separate category *c_B_* from *c_A_*. A problem here is that positive classification continues beyond the reasonable support of *c_B_*, including a third category (shown in blue). The second panel (b) shows a Gaussian distribution fit as a generative model of that data and the resulting conditional decision function. Note the green points from data *x* labeled *C_B_* have a cluster near the data labeled *C_A_* and other points farther away — because the region is wide, it yields weaker discrimination. The conditional decision function near *C_A_* and *C_B_* is impacted by data far from that decision boundary. The alternative proposed in this work, panel (c), uses statistical Extreme Value Theory (EVT) to estimate the probability distributions. With EVT, estimates are made separately for the tails at each decision boundary for a category, e.g., one for the lower decision boundary 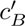, and another for the upper decision boundary 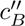.

The rest of this paper describes two human behavioral studies that collect category labels which are assigned to different types of stimuli under different configurations of reference stimuli. Generative EVT and Gaussian models are then compared to the human behavioral data to assess fidelity. In these experiments we make use of a gap-filling paradigm [Cohen et al., 2001, Hsu and Griffiths, 2010] for probing the representations underlying category learning, whereby a gap is left along the stimulus axis between a set of reference stimuli, and the subjects’ categorization of stimuli in that gap are evaluated in a two-alternative forced choice task. The extent to which Category A or Category B captures data points in the gap region provides important clues as to the underlying representation used by the subjects to categorize the stimuli. We conduct two experiments with different forms of stimuli. In Experiment 1, line segment stimuli that vary in a single dimension of length [Hsu and Griffiths, 2010] are used. In Experiment 2, more complex 2D rendered face sequences drawn from morphspaces [Folstein et al., 2012] are used as stimuli. In the rest of this introduction, we briefly review the possibilities for representation in category learning and previous behvorial studies making use of morph sequences. Then, in the following sections, we introduce our experimental methodology and results, as well as some discussion on the implication of these findings.

### Category Learning: Averages or Extrema?

Neuroscientists and psychologists have long studied how the human brain perceives and interprets the visual world, contributing to a deeper understanding of how sensory information is encoded. Over the past several decades, a prominent theoretical framework has emerged, suggesting that the brain adaptively models the statistical structure of visual inputs to extract task-relevant information. This line of research has led to influential developments in sparse coding [Olshausen and Field, 1996, Vinje and Gallant, 2000, Olshausen and Field, 2004, Foldiak and Endres 2008, Quian Quiroga and Kreiman, 2010, Chalk et al., 2018], probabilistic interpretations of perception [Berkes et al., 2011, Jern and Kemp, 2012, Gallistel et al., 2014, Block, 2018, Byrne, 2022], and related domains.

In psychology and cognitive science, modeling sensory input data shaped by human perception often begins with the use of central tendency models based on Gaussian distributions. For instance, [Tenenbaum and Griffiths, 2001] extend Shepard’s universal law of generalization by embedding it in a Bayesian framework, allowing generalization from multiple stimuli with arbitrary structure. That approach unifies spatial and set-theoretic models of similarity, reinforcing the idea that generalization often follows Gaussian-like decay in psychological space. [Minda and Smith, 2001] demonstrate that prototype models — based on central tendencies — can outperform exemplar models in categorization tasks involving complex or high-dimensional stimuli, reinforcing the value of Gaussian-like representations in perceptual modeling. [Feldman et al., 2009] explain the perceptual magnet effect using a Bayesian model in which perception is optimally biased toward category means, showing how central tendency shapes sensory judgments under uncertainty. [Briscoe and Feldman, 2011] show that prototype and exemplar models lie on opposite ends of the bias–variance spectrum, with human learners typically adopting an intermediate strategy, suggesting that central tendency (Gaussian-like) models capture only part of human categorization behavior. And most recently, [Battleday et al., 2020] show that combining rich visual representations with cognitive models reveals that simple prototype models — often based on central tendency assumptions — can effectively account for human categorization of natural images, performing comparably to more complex exemplar models.

However, vision science has surfaced much evidence supporting an alternative modeling approach that emphasizes the extremes of a distribution [Leopold et al., 2006a, Tsao and Livingstone, 2008, Freiwald et al., 2009, Groen et al., 2012b, Leopold et al., 2001a, Scholte et al., 2009, Tanaka et al., 2012]. Even though these findings suggest that emphasizing distributional extremes may be a biologically plausible strategy, the application of EVT to model sensory input and category boundaries remains underdeveloped in psychology and neuroscience. With respect to visual recognition, EVT modeling work has been more common in the field of computer vision.

[Scheirer et al., 2010] introduced the w-score, a robust score normalization method for recognition systems, based on EVT modeling of top non-match scores without assuming specific match/non-match distributions. Following that work, [Scheirer et al., 2011] proposed a method for predicting recognition performance using EVT-based post-recognition score analysis, and demonstrated its effectiveness across multiple recognition tasks using a Weibull-based statistical predictor. [Scheirer et al., 2012] used EVT techniques to calibrate and normalize multi-attribute spaces for attribute fusion and similarity search. [Jain et al., 2014] addressed open set recognition by modeling decision boundaries using EVT and introduced the *P_I_*-SVM algorithm to estimate the unnormalized posterior probability of category inclusion based solely on positive training data, with subsequent work modeling both the probability of category inclusion and exclusion [Scheirer et al., 2014]. Inspired by this idea, [Rudd et al., 2018] introduced the Extreme Value Machine (EVM), a theoretically grounded classifier based on EVT that enables incremental learning and open set recognition without relying on kernels, addressing limitations in traditional classifiers when handling unseen categories. More recently, [Vignotto and Engelke, 2020] examined the limitations of the EVM in certain recognition scenarios and introduced two algorithms that use EVT without using the geometry expressed by the known categories, which leverage the distance of test points from training categories.

### Human Behavioral Studies Using Morph Sequences

Morph sequences are used in behavioral studies to investigate how subjects perceive gradual transitions between stimuli, revealing the processes behind recognition, categorization, and perceptual continuity. By analyzing responses to subtle feature changes, researchers gain insight into identity boundaries and perceptual thresholds, advancing our understanding of how the brain processes complex, evolving visual input. Recent work has also touched on how decision boundaries emerge in recognition tasks, revealing how individuals resolve ambiguity when stimuli fall between well-defined categories. Faces are of particular interest to the research presented in this paper.

[Busey, 1998] examined how morphed faces relate to a psychological face space using multidimensional scaling. While morphs appeared more typical (or average) than their parent faces, consistent with geometric models, systematic biases such as age and adiposity effects suggest that morphing techniques can distort perceived similarity. [Schweinberger et al., 1999] explored how humans make categorization decisions when processing facial identity and emotion. Findings revealed an asymmetric influence: identity decisions were stable despite changes in expression, while emotion-based decisions were biased by identity — suggesting that identity exerts a stronger influence on perceptual decision-making than emotion. [DeBruine, 2004] extended previous research by using more realistic morphing techniques to manipulate facial resemblance between participants and child faces and showed that using morphs to increase facial resemblance boosted attractiveness and investment ratings, with no difference between men and women. [Busey and Arici, 2009] used morphed face stimuli in a forced-choice recognition task to show that participants were equally likely to choose studied and unstudied morphs but were more confident when selecting studied ones. This dissociation between recognition and confidence challenges Gaussian signal detection models and supports a sampling-based decision process that engages memory for individual items. [Blunden et al., 2015] investigated whether face morph dimensions are processed integrally or separably by analyzing response times in a categorization task. Results revealed individual differences, with some participants showing coactive (integral) processing and others using parallel, self-terminating (separable) strategies. More recently, [Ma et al., 2022] found that morphed multiracial faces are more likely to be classified as multiracial than real ones, highlighting that morph-based stimuli may not fully capture the perceptual judgments elicited by real multiracial faces.

This previous work on face morphs has explored perceptual, cognitive, and social decision-making, using morphs to study face space, feature integration, and categorization. While useful for controlled stimulus generation, these studies have not systematically examined how sampling methods influence decision-making or which models best capture behavioral changes across morphing regimes.

## Methods

### The Statistical Extreme Value Theory for Recognition

[Gumbel, 1954] demonstrated that for any continuous and invertible parent distribution, only three types of EVT distributions are needed to characterize the distribution of extrema. These depend on whether one is modeling maxima or minima, and whether the support is bounded above or below. These three canonical forms can be unified under the Generalized Extreme Value (GEV) distribution, defined as:

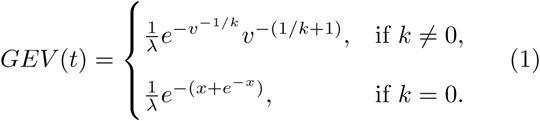

where *x* = *t - τ*, *v* = 1 + *k*(*t - τ*)*/λ*, and the parameters *k*, *λ*, and *τ* represent the shape, scale, and location, respectively. Varying the shape parameter *k* yields the three classical EVT distributions: *k* = 0 corresponds to the Gumbel (Type I), *k >* 0 to the Fŕechet (Type II), and *k <* 0 to the Reversed Weibull (Type III) distribution. The Gumbel distribution applies to data with unbounded support; the Frechet distribution applies when the support is bounded from below but unbounded from above (e.g., response times, minimal distances); and the Reversed Weibull applies when the data are bounded from above (e.g., maximal similarity). For cases where data are bounded from below, the standard Weibull distribution is also employed. In practice, many perceptual tasks involve bounded input domains, requiring combinations of models of maxima and minima, i.e., mixtures of Reversed Weibull and Weibull distributions.

The key insight from EVT [NIST, 2008] is that regardless of the form of the parent distribution, the distribution of extreme samples near decision boundaries will *always* converge to a member of the EVT family. That is, there is no need to assume a particular parametric form for the overall data distribution — whether it is a truncated binomial, a mixture of Gaussians, a log-normalized distribution, or even a bounded, multi-modal form [Bertin and Clusel, 2006], because the distribution of extremes near the boundary is theoretically guaranteed to follow an EVT distribution. While standard EVT assumes i.i.d. samples, this framework can be generalized to the configuration of exchangeable random variables [Berman, 1962], further extending its applicability to perceptual modeling.

With this key insight in mind, we use EVT to develop two models, the Probability of Inclusion (*τ* = positive) and Probability of Exclusion (*ε* = negative), and then combine them into a *P_τε_* model [Scheirer et al., 2014], which is the product seeking the probability that something is from the the first category and not from the second category. Finally, we can combine *P_τε_*with prior category probabilities to get the overall (unnormalized) posterior *P_π_*. To start, assume a hypothesis psychometric similarity *s_τ_*(*x, c*) such that a sample *x* is included in category *c*, and a hypothesis score *s_ε_*(*x, c*) such that *x* should be excluded from *c*, where we assume larger scores imply more likely inclusion/exclusion.

While it is straightforward to develop the decision model for any EVT distribution, for simplicity of presentation let us assume the inclusion scores are bounded from below and the exclusion scores bounded above, which dictates the distribution for the extremes to be a Weibull and Reversed Weibull respectively. Note in both cases we are using a few extreme values to fit the long-tail of the distribution, not its bounded half.

To estimate the probability for any psychometric score *x* belonging to category *c*, we use the appropriate cumulative distribution function with the associated parameters. Given *x*, we have two independent estimates: one for inclusion, *P_τ_*, based on the Weibull CDF derived from the match data for category *c*, and another for exclusion, *P_ε_*, based on the Reversed Weibull CDF from the non-match estimates for category *c*. This yields our category configurational density estimation, the Probability Inclusion Exclusion estimate *P_τε_*, defined by the product

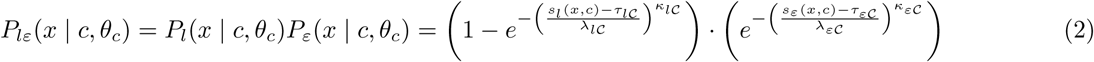

where θ*_c_* represents the parameters for an EVT fit. For a given stimulus, one must find the nearby extrema from the different categories, compute *P_τε_*, and then predict the category with the maximum probability.

### Gap Generalization with Varying Exposure Distributions

If we consider any computational model of learning, we can compare models by considering their predictions. Given that extreme value representations are naturally focused on extremes, paradigms that require subjects to generalize beyond previous experience present a powerful opportunity to expose differences between EVT and prior models. Along these lines, Hsu and Griffiths [Hsu and Griffiths, 2010] used an elegant paradigm (adapted from previous studies [Cohen et al., 2001, Rips, 1989, Smith and Sloman, 1994, Stewart and Chater, 2002]) for probing the representations underlying category learning. To explain the core concept, we start by extending this gap filling paradigm, with the original concept depicted in Fig. 2. This gap paradigm is powerful; however, because experiments cannot directly measure the psychometric axis, the differences in the gap are a mixture of both normalization of the psychometric space and the response model for categorization [Shepard, 1980, Smith, 2006, Navarro, 2007].

**Figure 2:**
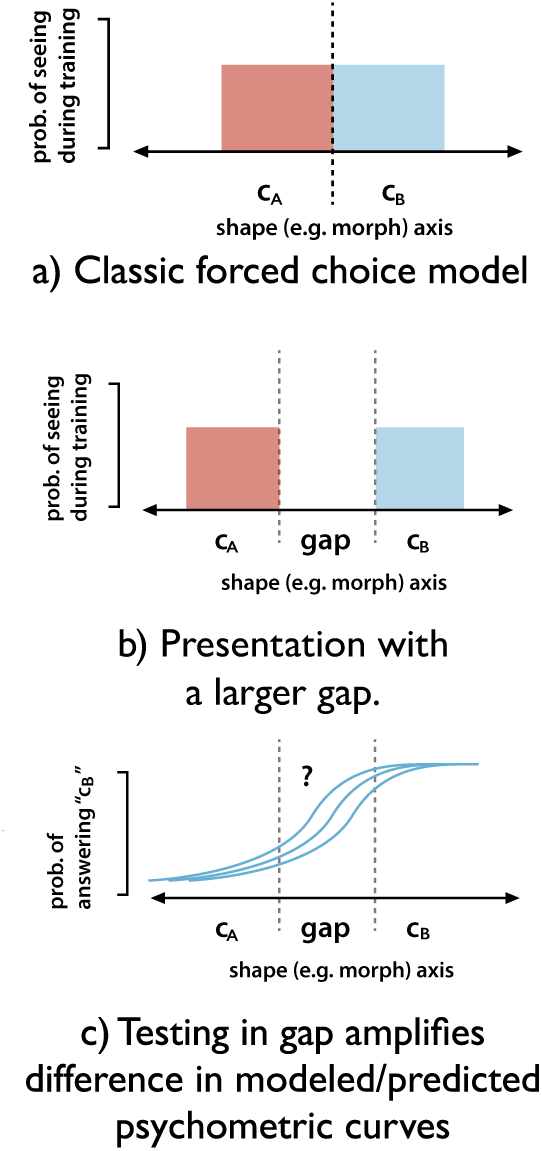
Conceptual framework for the Gap-filling paradigm, designed to assess how different models generalize in unseen regions of the stimulus space. Panel (a) shows a typical forced-choice training setup with two categories, *c_A_* and *c_B_*, defined over adjacent stimulus regions. Panel (b) introduces a gap between the categories, leaving the intermediate region unobserved. Panel (c) illustrates the key consequence of this design: when testing stimuli within the gap, model predictions diverge substantially. The resulting psychometric curves reveal the differing assumptions of each model regarding generalization beyond the observed data.

The key feature of extreme value models is that they are fit using data that lie in the tails of the distribution. This property gives extreme value distributions their well-known ability to accurately estimate rare events, but it also makes them relatively insensitive to data in the middle of the distribution to be modeled. For example, since Gaussian models, in contrast, are affected by data across their range, every data point contributes to the mean and variance of the distribution, which govern the Gaussian shape. This difference provides experimental leverage to distinguish between these two possible models. In the experimental designs that follow, subjects are exposed to different distributions of reference stimuli in an unfamiliar stimulus space and then probe the resulting category organization.

In all experiments, subjects are first trained to perform a two-alternative forced choice task along one axis of a stimulus space. In a variation on the experiments of Cohen et al. and Hsu and Griffiths, a gap in training exposure is left along the stimulus axis in the reference stimuli shown to a subject, and the categorization of stimuli in that gap is the task performed by a subject. The degree to which Category A or Category B “captures” data points in the gap region provides important clues as to the underlying representation used by subjects to categorize the stimuli.

The experiments start with a symmetric case, where training examples are drawn from distributions for each category such that both categories have equal opportunity to capture gap stimuli (though response biases could still exist), designed such that the Gaussian and EVT models should behave in a similar manner. In general, we cannot know how the units of our designed stimulus axes map onto the units of the subjects’ underlying perceptual representations, and thus this “symmetric” case represents an important control against which to compare experimental configurations in which training distributions are manipulated for each category.

As shown in Figure 3, EVT and Gaussian model predictions can diverge strongly as the distribution of reference stimuli changes. As examples are redistributed for one of the categories (e.g., by making the tails of the sampling distribution longer or “fatter”), the Gaussian and EVT models make substantially different predictions, since the EVT model fits the extrema and is relatively insensitive to data far from the category boundary, while the Gaussian model incorporates data across the whole category. In the enriched tail configuration, the shift in model predictions, relative to the uniform baseline, can depend on how the tail is sampled. Factors such as the number of enriched samples, the range of two tails, and their proximity to the category boundary all influence the outcome. For EVT, enriched sampling near the boundary increases local density and compresses the response function, often pulling the psychometric curve closer to the true midpoint of the gap and reducing bias. In contrast, Gaussian models may respond to the broader mass of enriched points, potentially increasing variance, which can lead to a larger bias away from the midpoint. In the long tail configuration, reference stimuli extend further from the category center without concentrating near the boundary. EVT predictions remain stable, as the model emphasizes extremes rather than distributional spread. Gaussian models integrate over the full range, leading to increased mean shifts that pull the psychometric curve further into the gap.

**Figure 3:**
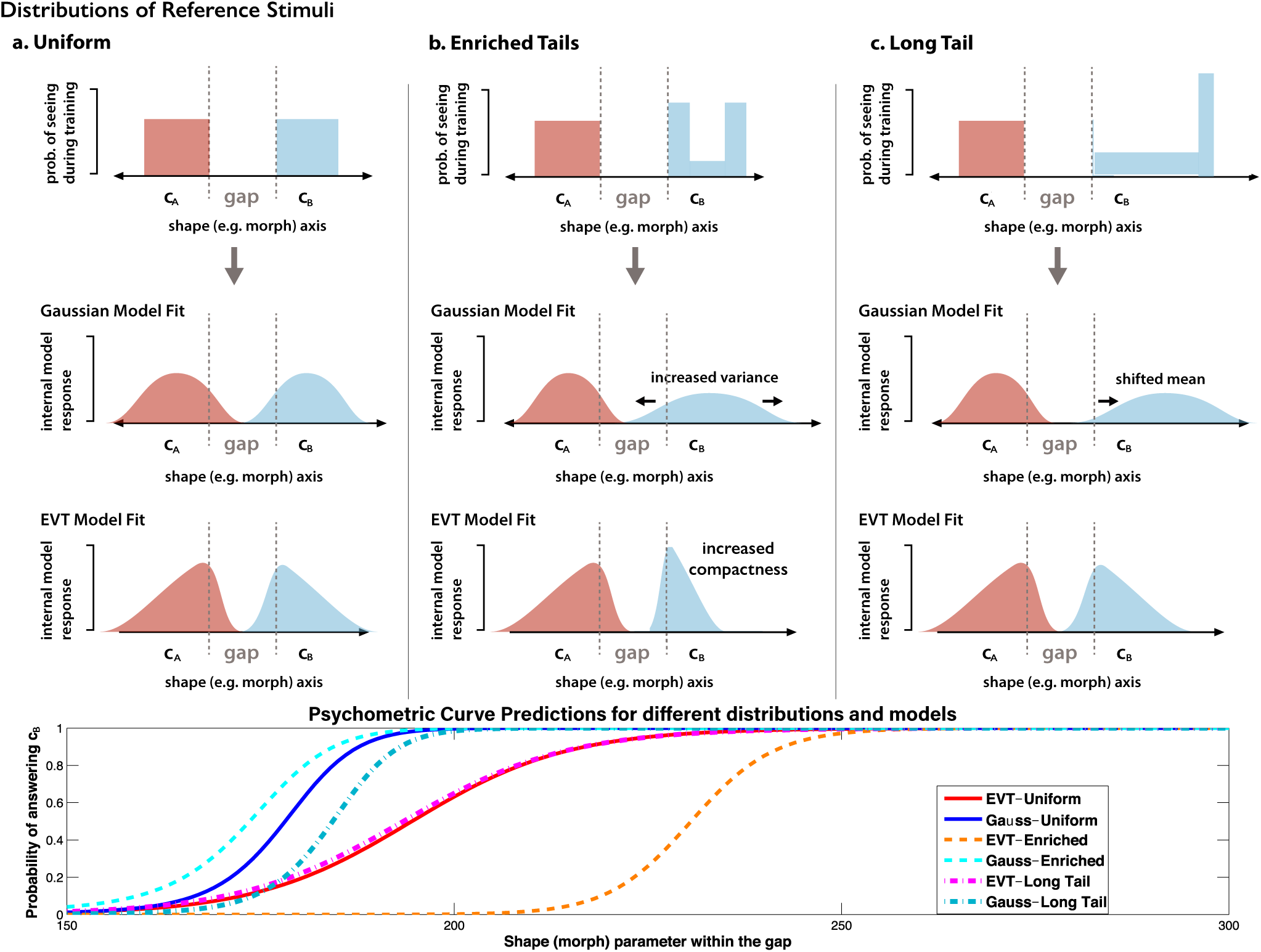
Distributional Variation Paradigm: Distinguishing Gaussian and EVT Predictions. A central component of the paradigm is the training of a two-category discrimination task along a parametric stimulus axis. first row: the probability of drawing examples from category A (*c_A_*) during training is shown in red, and the probability of drawing category B (*c_B_*) is shown in blue. In all configurations, a gap region where no training examples were shown is imposed between the two categories. The second row shows Gaussian model fit predictions for each training distribution, and the third row shows corresponding EVT model fits. Finally, the fourth row shows simulated psychometric predictions derived from these model fits; these curves are not drawn from real experimental data, but instead illustrate the predicted differences between Gaussian and EVT models under different distributions. This simulation highlights that even small changes in the reference distributions can lead to substantial divergence between models, particularly within the unseen gap region. Data used to generate these curves can be found in Section A in the Appendix of this paper.

For all configurations, the direction of the experimental axis in the stimulus space is varied across subjects to guard against unexpected stimulus effects (e.g., unintended perceptual non-uniformities in the stimulus space). In particular, the distributional manipulations described above will be counterbalanced across subjects, such that all manipulations on the “left” side of a particular axis for one subject will be mirrored on the opposite side of the axis for another. Experiments are conducted across two different stimulus spaces, again to reduce unexpected biases.

## Gap Generalization Experiments

### Experiment 1: Human Gap Generalization Modeling Considering Line Segments as Stimuli

For an initial study to analyze the applicability of the EVT-based modeling approach to recognition in humans, we adapted the design proposed by [Hsu and Griffiths, 2010], which used line segments as stimuli to investigate categorical perception. Similar to their approach, we employed line segments to construct our stimulus space and explored how category learning generalizes in untrained regions. In the original study, the training and testing phases were distinct: participants first viewed training stimuli, after which these stimuli were removed, and only then were they asked to categorize new test stimuli. Different from this, our experiment provided continuous visual access to a set of reference stimuli during the testing phase. This adjustment simplified the task for participants and resulted in smoother, less noisy behavioral responses, while preserving the essential structure of category generalization in gap regions.

In this experiment, subjects were presented with a series of line segments of varying lengths that were grouped into two categories: short lines (Category A) and long lines (Category B). The subjects were instructed to assign instances of line segments to each of these groups. Most importantly, a gap was left in the range of lengths, such that subjects were not shown line segments of intermediate length as reference stimuli, and the distribution of lengths in one category (the longer line segments) was varied.

Consistent with previous studies, Hsu and Griffiths found that humans are sensitive to category variability and were more likely to assign an object equidistant from the mean of two categories to the category with higher variance. This phenomenon of the higher variance category “capturing” previously unseen intermediate stimuli is consistent with the predictions of a Gaussian generative model — the higher variance Gaussian “spills over” into the gap.

Using similar experimental regime as Hsu and Griffiths, we led a study at the University of Notre Dame^1^ with 25 subjects for the three configurations shown in Figure 3: human generative (uniform), long tail, and enriched tail. The human results were then compared to EVT and Gaussian models, with an assumption that distributions emphasizing the extrema at the edges of the gap region might better explain subjects’ bias toward the high variance category.

#### Data and Experiment Design

In this study, we utilized horizontal line segments with varying pixel lengths, plotted against a uniform white background measuring 800 pixels in width and 300 pixels in height. For reference stimuli, we first generated five line segments for category A, which were always sampled uniformly, as shown in Figure 3. These line segments were shared across all three experimental configurations. For category B, we generated three sets of line segments, each set corresponded to a different sampling configuration: uniform, enriched tail, and long tail. Each set contained five line segments, sampled according to the respective distribution. In all images, the segments were evenly spaced across the canvas to maintain a consistent and structured layout.

Additionally, we generated eight testing line segments, each of which was centrally positioned on an identical 800 × 300 white background. This design maintained consistency across all images while facilitating a controlled comparison between categories.

As shown in Figure **??**, for the reference stimuli, line segments in category A had lengths ranging from 110 to 150 pixels, while those in category B ranged from 300 to 600 pixels. For the testing stimuli, we selected line segments with lengths between 167 and 283 pixels. This range was chosen to ensure a clear separation between the two reference categories while avoiding overlap with either category A or B. As mentioned, we had three different sampling configurations when it came to picking reference stimuli for category B. Table 1 shows the lengths of reference line segments used for our study. As for the test images, we selected eight fixed line segments for all three setups: 167, 183, 200, 216, 233, 250, 267, 283. With this design, we had 24 questions in our survey in total (three sampling configurations and eight test line segments/questions for each configuration).

**Table 1:** Reference line segment pixel length for category A and category B for the three sampling configurations.

| Configuration | Reference Line Segments<br>for Category A | Reference Line Segments<br>for Category B |
| --- | --- | --- |
| Uniform | 110, 120, 130, 140, 150 | 300, 375, 450, 525, 600 |
| Long Tail | 110, 120, 130, 140, 150 | 560, 570, 580, 590, 600 |
| Enriched Tail | 110, 120, 130, 140, 150 | 300, 325, 350, 575, 600 |

#### Web Application for Behavioral Experiments

To facilitate the experiments, we developed an online data collection web application with Python and Flask. This application ran on a server under our control, and was accessed by subjects participating in our experiments via their own web browsers. The full process of the data collection is included in Section B in the Appendix of this paper.

The study began with some introductory text outlining the background of the experiment, followed by a consent page detailing user privacy and terms of participation. Once participants agreed to the terms and configurations, they were directed to an instruction page. This page provided an overview of the experimental interface, explaining the purpose of each component and guiding participants on how to make a selection for the questions. Participants were then directed to a practice page where they answered a sample question to familiarize themselves with the process of selecting an answer. Upon completing the practice task, the website transitioned to the main survey.

Figure 4 shows an example question from the main survey. Each question asked a subject to decide which category a test line segment belonged to. For each question, three rows of line segments were shown. In the first row, five line segments were shown in a green box, and they were considered to belong to the same Category A. In the second row, another five line segments were shown in a red box, and they were considered to belong to the same Category B. In the third row, one line segment was shown, and the subject needed to decide whether the line segment in it belonged to Category A or Category B. In the last row, two options were shown. The participant could select the first option if they thought the line segment in the third row was from Category A, or select the second option if they thought the line segment in the third row was from Category B. The participant could click the (1) or (2) buttons with their cursor or press the (1)–(2) keys on their keyboard to make their selection. After a participant finished all of the questions, a survey code appeared on the screen and they were asked to submit the code to the research assistant conducting the experiment.

**Figure 4:**
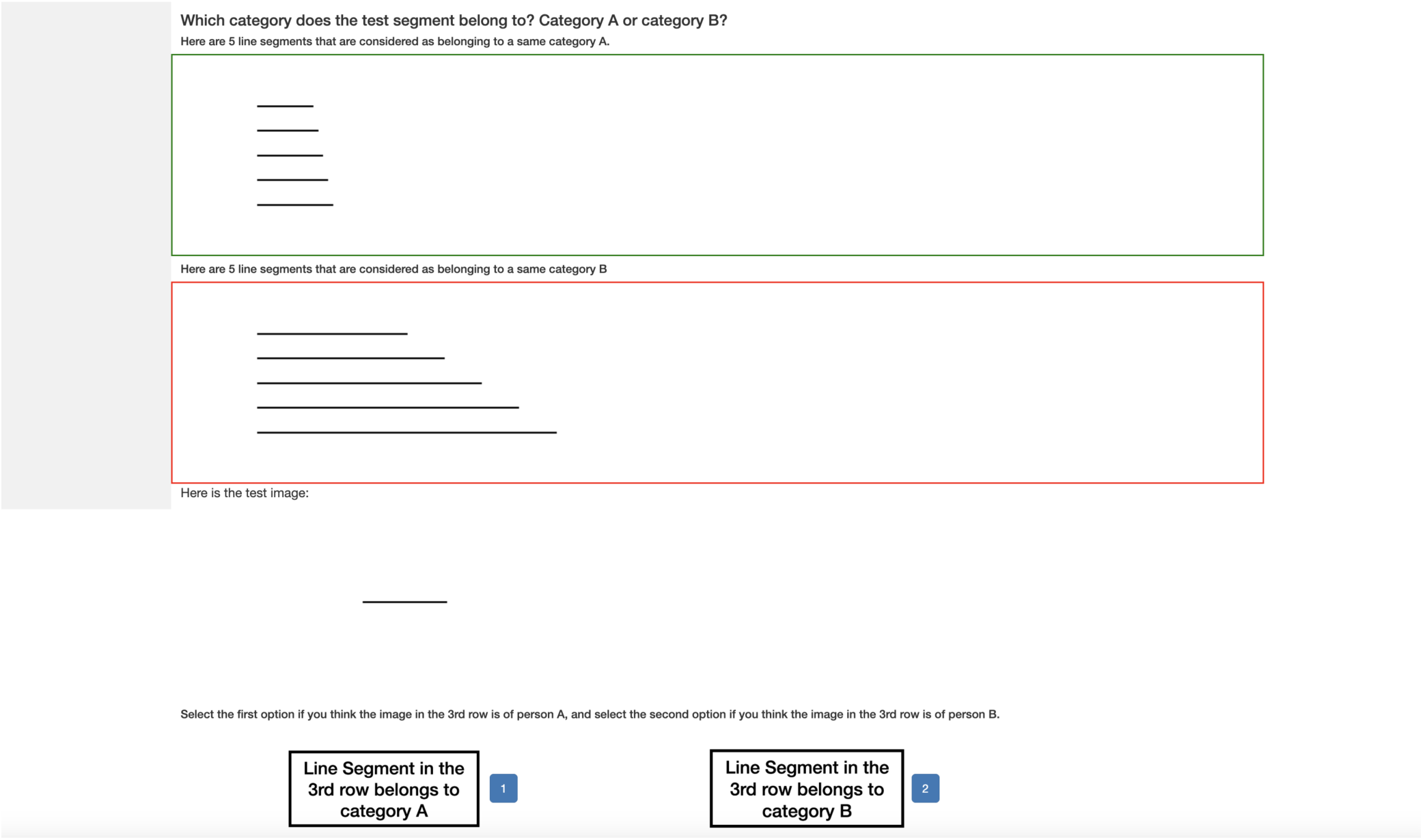
A question from the main survey. For each question, three rows of line segments were shown. In the first row, five line segments were displayed in a green box and were considered as belonging to Category A. In the second row, another five line segments were displayed in a red box and were considered as belonging to Category B. In the third row, one line segment was shown, and the participants needed to decide whether the line segment belonged to Category A or Category B.

#### Statistical Modeling

To evaluate how different computational models generalized under varying reference stimuli distributions, as well as how their behavior compared to human subjects, we considered two modeling frameworks: a Gaussian-based Naive Bayes classifier and an EVT-based model. Both models were tested using the same fixed set of eight transfer stimuli, with lengths spanning the unseen gap region.

The Gaussian model assumed that category membership could be modeled using normal distributions estimated from the full set of training examples in each category. We followed the algorithm of Gaussian Naive Bayes to calculate the probabilities. In this experiment, a separate Gaussian distribution was fit to each category’s reference stimuli, utilizing the Python scikit-learn method `sklearn.naive_bayes.GaussianNB.fit()`. Reference stinuli for Category A remained fixed across all configurations, while the training stimuli for Category B varied depending on the sampling configuration, as shown in Table 1. To fit the model, a Gaussian distribution was estimated for each category independently by calculating the mean and standard deviation of the reference stimuli. Given a test stimulus, the model evaluated the likelihood of that stimulus under each category’s distribution using the Gaussian cumulative distribution function (CDF). Assuming equal priors, the posterior probability for each class was then computed via Bayes’ Rule, with `sklearn.naive_bayes.GaussianNB.predict()`. The predicted category for each stimulus was determined by choosing the one with the higher posterior probability.

The EVT model offered an alternative to Gaussian modeling by focusing not on the full distribution of training data, but instead on the extremes — the most diagnostic values near the decision boundary. In this framework, we estimated category membership using two probability components: the Probability of Exclusion (PoE) from Category A, and the Probability of Inclusion (PoI) from Category B. For each sampling configuration (uniform, enriched tail, long tail), the EVT model also utilized the training data (segment lengths) described in Table 1. We first fit EVT models with a tail size of three, which was the minimum number of data points required to fit an EVT model. Given that the reference stimuli for each category contained only five samples, selecting three tail points struck a balance between statistical validity and practical constraint. To compute PoE, we used the three largest values from Category A’s training data to fit a reversed Weibull distribution (using the `weibull_max` method in the Python SciPy `scipy.stats` module), capturing the tail behavior as data diverged from Category A. Likewise, PoI was modeled using a Weibull distribution (`weibull_min` in the `scipy.stats` module) fit to the three smallest values from Category B, estimating the degree to which a stimulus resembled the boundary-extreme elements that belonged to Category B. This gave us estimates of shape, scale, and location parameters for PoI and PoE, respectively. With these parameters, we generated PoI and PoE for each testing length, and the final probability of a testing point was computed by multiplying the PoI and PoE of that point given the estimated parameters.

### Line Segment Experiment Results

#### Human vs. EVT vs. Gaussian

Figure 5 compares the categorization performance of human participants, EVT models, and Gaussian models for the line segment experiment under three different configurations. Human participants show clear shifts in generalization curves depending on the training configuration: enriched tails lead to lower probabilities of choosing Category B across the gap, comparing to the uniform configuration, while long tail distributions substantially suppress the likelihood of choosing B. The EVT models closely follow these human patterns, reflecting both the downward shift in the enriched tail configuration and the significant downward shift in the long tail configuration. In contrast, the Gaussian models exhibit very limited capability to match human curve shifts. For both uniform and enriched tail configurations, the Gaussian model quickly saturates to near-certain predictions of Category B. More importantly, the enriched tail curve shifts in the opposite direction compared to the human curve shift. As for the long tail configuration, it consistently predicts Category A across all test stimuli. These results highlight the limitations of Gaussian-based modeling in handling distributional changes and underscore the advantages of EVT-based modeling, which more closely follows the human behavioral responses.

**Figure 5:**
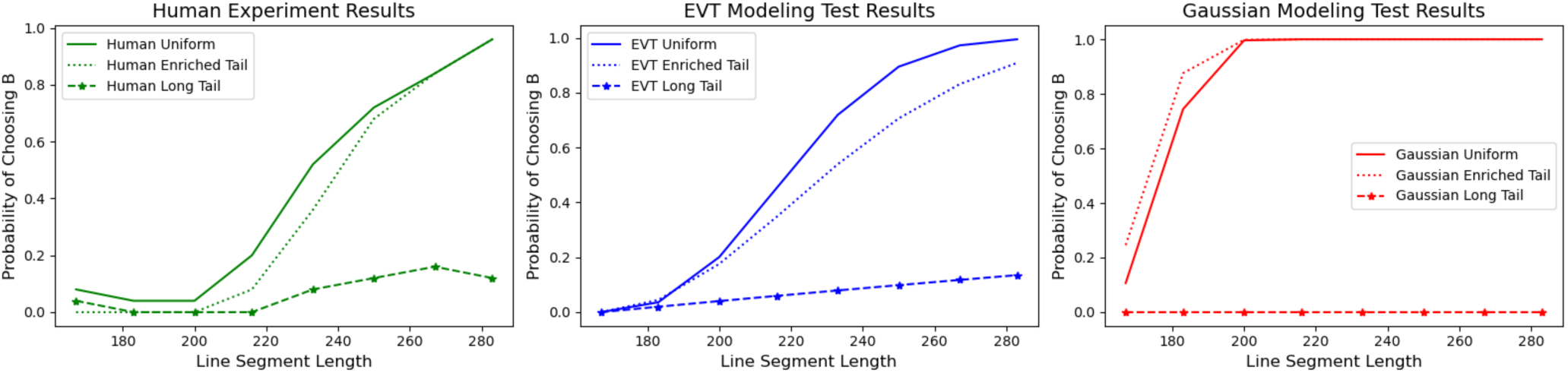
Comparison of categorization results across human participants, EVT-based models, and Gaussian models. Each subplot shows the probability of categorizing test stimuli into Category B, plotted against line segment length. The left panel displays human responses under three configurations: uniform, enriched tail and long tail. Human responses shift appropriately under the enriched tail configuration and show limited generalization under the long tail configuration, indicating sensitivity to distributional changes. The center panel presents results from EVT-based models, which successfully reproduce both the downward shift in the enriched tail configuration and the attenuated choice probability in the long tail configurations, closely matching human behavior. In contrast, the right panel shows Gaussian model predictions, which fail to capture the long tail suppression and instead produce overly confident categorization, particularly under the uniform and enriched configurations. These results suggest that EVT-based models better reflect human sensitivity to different tail configurations in category learning.

#### Statistical tests for normality

To evaluate whether human categorization aligns with Gaussian assumptions, we conducted the Shapiro–Wilk test [Shapiro and Wilk, 1965], a standard method for assessing normality in small to moderate samples. This test returns a p-value indicating the likelihood that a dataset is normally distributed. We applied this test to human responses for each of the eight test segment lengths, using `scipy.stats.shapiro`. Each segment had 25 binary responses (Category B or not), and we tested these separately under all three configurations (uniform, enriched tail, long tail). Across all segments and configurations, p-values were consistently very low (e.g., 10^−6^ or smaller), indicating strong deviation from normality. This result underscores a key limitation of Gaussian-based models: their assumption of symmetric, bell-shaped distributions. The clear non-normality in responses suggests that Gaussian models are ill-suited to this task and supports the use of alternatives like EVT, which better capture skewed or heavy-tailed patterns.

We further assessed distributional shape using the Kolmogorov–Smirnov (K–S) test [Massey, 1951], a non-parametric method that compares the empirical cumulative distribution function (ECDF) to that of a reference distribution. Here, we tested whether binary responses for each segment could plausibly come from a normal distribution with matching mean and standard deviation using `scipy.stats.kstest`. All p-values were well below 0.05 (ranging from 10^−3^ to 10^−7^), confirming significant deviations from normality. These findings corroborate the Shapiro–Wilk results and further justify the use of non-Gaussian models such as EVT for capturing human categorization behavior.

### Experiment 2: Human Gap Generalization Modeling Considering Face Morph Sequences as Stimuli

Building on the foundation of the line segment experiment, we extended the setup to more complex and natural stimuli: human face morph sequences. This progression from simple to natural stimuli allows us to assess realistic visual appearances where more ambiguity is present, compared to the line segments.

#### Human Face Dataset Description

To collect data for the study, we used the “Labeled Faces in the Wild (LFW)” dataset. The LFW dataset was chosen because it provides a large collection of real-world, unconstrained face images with identity labels, making it well-suited for studying perceptual categorization under naturalistic variability. In addition to variation in pose and expression, this dataset also includes substantial diversity in race, gender, background, and lighting configurations. The LFW dataset contains more than 13,000 images of faces of public figures collected from the web. Each face has been labeled with the name of the person pictured. 1,680 of the people pictured have two or more distinct photos in the dataset. It was collected by the University of Massachusetts Amherst [Huang et al., 2007b]. There are four different sets of LFW images including the original set and three different types of “aligned” images. The aligned images include “funneled images” [Huang et al., 2007a], LFW-a, which uses an unpublished method of alignment, and “deep funneled” images [Huang et al., 2012]. Among these, LFW-a and the deep funneled images produce superior results for most face verification algorithms over the original images and over the funneled images. For our experiments, we utilize the deep funneled version, which will be referred to as “LFW deep funneled” later in this paper. This dataset contains 13,233 images for 5,749 people.

#### Human Face Data Selection

To ensure the quality of the face morph sequence, we carefully selected images based on a few key factors including gender, skin color, face angle, and the clarity of the face in the image. For skin color, we categorized the images into dark and light tones, while gender was classified as male or female. Face angle was also considered, with images representing left, right, and front perspectives. Additionally, we ensured that the selected images featured clearly visible faces, free from any occlusions such as hair, accessories, or other objects or shadows. These criteria were chosen to maintain consistency across the sequences and to enhance the accuracy and reliability of the resulting morphs. By controlling for these variables, we aimed to create a dataset that is both diverse and representative, while also ensuring high visual fidelity for human observers. The exact number of images selected for each key factor can be found in Section C in the Appendix.

#### Face Morph Sequence Generation

In order to generate human face morph sequences, we utilized an open-source project called “face morpher^2^” (hereafter referred to as “the morpher”). The morpher, implemented with Python and OpenCV [Bradski, 2000], morphed one face into another by leveraging facial landmark detection and fusion. The process involved taking a 500×500 image, cropping the face region, and removing the background to focus solely on the facial features. Facial landmarks were then detected using the dlib 68 facial landmark detector [Kazemi and Sullivan, 2014] in both the source (Person A) and destination (Person B) images to identify key points such as the eyes, nose, and mouth. The images were aligned to a standard size, and intermediate facial landmarks were calculated for each frame of the morphing sequence by interpolating between the source and destination landmarks. Using these landmarks, the software applied affine transformations to warp the facial regions of the source and destination images. The warped images were blended together, with the blending weights gradually shifting from the source to the destination face. The resulting frames were saved as individual images as well as compiled into a video, creating a smooth and visually seamless morphing effect.

For each morph pair, we randomly selected one person A and one person B, but also ensured they had the same skin tone, gender, and face angle before choosing one image of each. After that, the morpher produced *n* frames that morphed person A to person B. To make sure we had detailed morphs and the necessary amount of data for the experiments, we picked *n* = 140 in this study, with each sequence containing 142 images, including person A and person B. The resulting images were saved during the process.

#### Human Response Data Collection Application

In order to collect data with human subjects, we developed a second data collection web application with Python and Flask. This application is similar to what we used for the line segment experiments, with subjects again using their own web browsers, but adapted to the face morph data. The study also began with introductory text outlining the background of the experiment, followed by a consent page, an instruction page, and a practice task. After the practice task was completed, the web application transitioned to the main survey.

The question asked a subject to decide which person a test image was more similar to. The subjects were instructed to try their best to answer the questions, taking as much time as needed. For each question, three rows of images were shown. In the first row, five images were displayed in a green box and were considered as belonging to the same person A. In the second row, another five images were displayed in a red box and were considered as belonging to the same person B. In the third row, one image was shown, and the participant needed to decide whether it was of person A or person B. In the last row, two options were shown. The participant was instructed to select the first option if they thought the image in the third row was of person A, and the second option if they thought it was of person B. The participant could click the (1) or (2) buttons with their cursor or press (1) or (2) on their keyboard to make their selection. After a participant finished all the questions, a survey code appeared on the screen, and they were asked to submit the code to the research assistant conducting the experiment. Full data collection process details can be found in Section D in the Appendix of this paper.

#### Experiment Design

For each face morph sequence, we generated 140 frames of morph images. This meant that in each sequence, frame 1 was the image of person A, frame 142 was the image of person B, and all 140 frames in between were combinations of A and B. We chose frames 1 to 8 as the range of Category A, and frames 83 to 142 as the range of Category B. This design carefully mirrored the structure used in Experiment 1. In the line segment experiment, Category A covered a narrow range (110–150, range = 40), while Category B spanned a much broader interval (300–600, range = 300), making B approximately 7.5 times wider than A. Following the same ratio, we selected frames 1–8 as the reference range for Category A in the face morph experiment, and frames 83–142 for Category B. This preserved the 7.5:1 ratio in range, ensuring the statistical asymmetry between categories was comparable across both line segment and face morphs experiments. To sample Category B for the long tail and enriched tail configurations, we selected a tail size of five. This meant that for the long tail experiment, frames 138 to 142 were considered the tail. We sampled the tail with a weight of 0.9 and the other frames with a weight of 0.1. For the enriched tail experiment, frames 83 to 87 were considered the head of Category B, and frames 138 to 142 were considered the tail of Category B; head and tail were each sampled with a weight of 0.45. All other frames, except for the head and tail, were sampled with a weight of 0.1. Following this design, we obtained reference frames for all three configurations, as shown in Table 2.

**Table 2:**
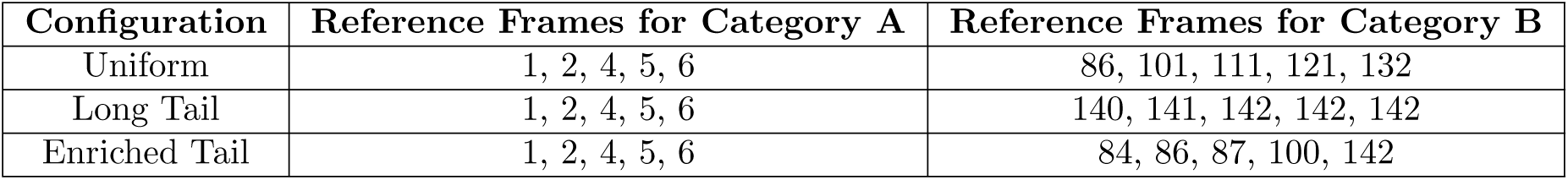
Reference frame indices for Category A and Category B for 3 experiments.

| Configuration | Reference Frames for Category A | Reference Frames for Category B |
| --- | --- | --- |
| Uniform | 1, 2, 4, 5, 6 | 86, 101, 111, 121, 132 |
| Long Tail | 1, 2, 4, 5, 6 | 140, 141, 142, 142, 142 |
| Enriched Tail | 1, 2, 4, 5, 6 | 84, 86, 87, 100, 142 |

As for the test images, we selected 20 fixed frames for testing across all morph sequences and all three configurations: 36, 38, 40, 42, 45, 47, 50, 52, 55, 57, 60, 62, 65, 67, 70, 72, 74, 76, 78, and 80. This test range was chosen to lie entirely within the unseen gap between Categories A (frames 1–8) and B (frames 83–142), analogous to the design in Experiment 1, where test stimuli were positioned in the perceptual gap between narrowly distributed A samples and broadly distributed B samples. The selected frames provided dense, uniform coverage across the boundary region, allowing for fine-grained assessment of generalization behavior and decision boundary shifts under different reference stimuli distributions. This setup ensured comparability with Experiment 1 while also leveraging the continuous nature of the face morph space.

#### Human Response Data Collected

We generated 90 face morph sequences using 180 carefully selected face images, and divided them into five survey sets, each containing 18 morph sequences. For every morph, we selected 20 test frames within the unseen gap region, resulting in 360 questions per survey. To mirror the structure of Experiment 1, each survey was completed under three distinct configurations: uniform, enriched tail and long tail. We recruited 25 participants in total, assigning five subjects to each survey set. All participants completed the three configurations in a fixed order — uniform first, then enriched tail, followed by long tail — and were instructed to take as much time as needed. On average, it took a subject 60 to 70 minutes to complete all three configurations. This design ensured a balanced and comprehensive collection of human responses, yielding 27,000 data points overall (90 morphs × 20 questions × 5 participants × 3 configurations).

#### Human Face Data Pre-processing for Statistical Modeling

Because the EVT model required one-dimensional input, we processed and compressed the face morph images into one-dimensional data for each image. For each frame in the reference and testing sets, we computed the distance from the morph image to the corresponding image of person A and person B. To achieve this, we used a ResNet-50 model that is pre-trained with the VGG-Face dataset [Parkhi et al., 2015]. The VGG-Face dataset is a large-scale face recognition benchmark containing approximately 2.6 million images from 2,622 distinct identities. It was originally introduced alongside models based on the VGG-16 [Simonyan and Zisserman, 2015] architecture, which achieved strong performance on early face recognition tasks in computer vision. Building on this foundation, subsequent community efforts extended the use of the VGG-Face dataset to train deeper and more powerful architectures, such as ResNet-50 [He et al., 2016]. The VGG-Face ResNet-50 model employs the ResNet-50 architecture, a 50-layer deep convolutional neural network with residual connections that facilitate efficient optimization and stable training. Trained on the VGG-Face dataset, the VGG-Face ResNet-50 model extracts discriminative 2048-dimensional facial feature embeddings suitable for tasks such as face verification, clustering, and similarity analysis. Compared to earlier VGG-16 and VGG-19 [Simonyan and Zisserman, 2015] models, VGG-Face ResNet-50 offers improved recognition performance due to its deeper structure and enhanced learning capacity. The features for each image, including person A’s image (frame 1), person B’s image (frame 142) and all reference and testing frames, were extracted from this model. We then calculated the Euclidean distance from each image to person A’s and person B’s reference images, so that each frame has a distance score to Person A (dist_to_A) and a distance score to person B (dist_to_B). These distance values were used as the one-dimensional input for the EVT and Gaussian models in our analysis later.

#### Statistical Modeling

Similar to Experiment 1, we utilized Gaussian Naive Bayes to model the probabilities. In this experiment, the Gaussian model was trained globally across all surveys by aggregating reference stimuli as training data for the models. To align with Experiment 1, we aimed at calculating the probability of a point belong to Category B, and we used distance scores to A (dist_to_A) as the input for Gaussian model. Specifically, 450 training data points (5 frames x 90 surveys) were used for Category A and similarly 450 training data points were used for Category B. A separate Gaussian model was fit to each Category based on the aggregated training data, using the calculated euclidean distance scores as input features. During testing, 20 fixed test frames were evaluated. For each frame index, we utilized 90 corresponding test samples (one from each survey), computed the posterior probability for Category A and Category B for each sample under the fitted model based on Bayes Rule, and averaged the 90 probabilities to obtain the final probability score for that frame. This approach was repeated for all three configurations (uniform, long tail, enriched tail).

For the EVT model in Experiment 2, we first obtained distance scores for reference stimuli in Category A and Category B. Specifically, we had 450 distance scores for distance to A and 450 scores for distance to B for each category (5 frames × 90 surveys). For distance to A, the scores were sorted in ascending order, while for distance to B, the scores were sorted in descending order. We then defined a tail size of ratio 0.2, selecting the most extreme 90 scores (0.2 × 450) from each category for model fitting. Given that the distance scores are strictly positive and naturally bounded both below and above – never reaching zero or infinity – we modeled the probability of inclusion (PoI) and the probability of exclusion (PoE) separately. PoI was estimated by fitting a reversed Weibull distribution to the distance scores to Category B, reflecting the idea that smaller distances imply stronger inclusion in B. Conversely, PoE was estimated by fitting a standard Weibull distribution to the distance scores to Category A, with larger distances indicating stronger exclusion from A. These provide estimations of shape, scale and location parameters. During testing, for each test frame index, we used the parameters estimated above to evaluate all 90 corresponding test points (one from each survey), computed PoI and PoE for each point, and obtained the probability score by multiplying PoI and PoE. The final probability for each frame was then determined by averaging the 90 individual scores. This procedure was performed separately for each of the three configurations: uniform, enriched tail, and long tail.

### Human Face Morph Experiment Results

#### Human vs. EVT vs. Gaussian

Figure 6 illustrates the categorization results for the face morph experiment across human participants, EVT models and Gaussian models. In the human data (left panel), participants show strong and relatively smooth transitions from low to high probability of choosing Category B across the test frames. Enriched tail training slightly shifts the curve upward compared to the uniform baseline, while long tail training results in a noticeable suppression of B responses, particularly in the mid-gap region. The EVT models (middle panel) are able to capture these shifts in direction: enriched tail training elevates the psychometric curve relative to the uniform configuration, and long tail training substantially reduces the probability of choosing B. Although the absolute scale of EVT model predictions is somewhat lower than human responses. In contrast, the Gaussian models (right panel) predict near-certain assignment to Category B across all frames and configurations, failing to differentiate between training distributions. This pattern again highlights the Gaussian model’s insensitivity to training distribution manipulations, while the EVT model more effectively reflects the distributional effects observed in human generalization behavior.

**Figure 6:**
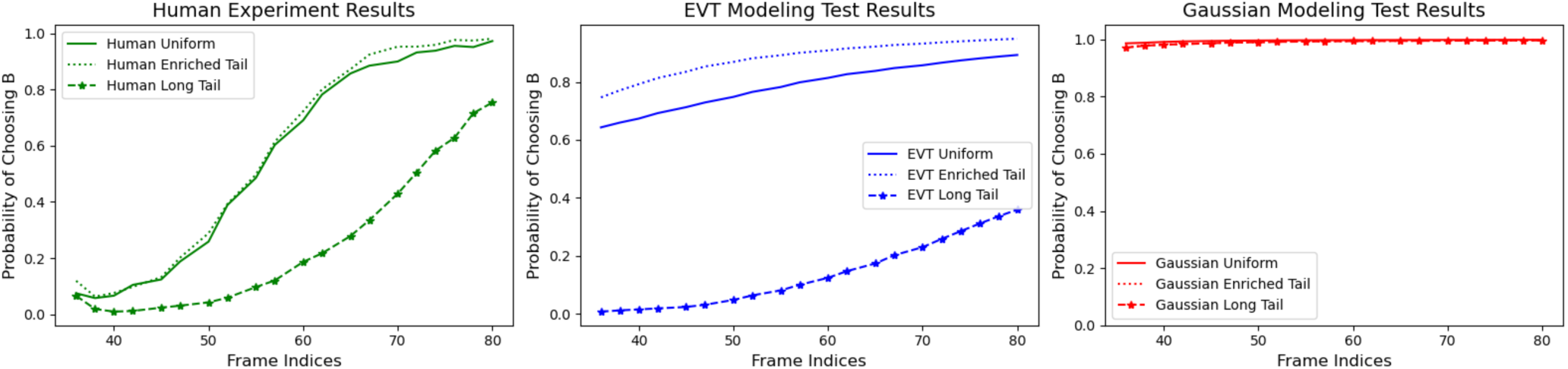
Comparison of categorization results across human participants, EVT models, and Gaussian models for Experiment 2 under three different configurations. Each subplot shows the probability of categorizing test frames as belonging to Category B, plotted against frame indices. The left panel displays human behavioral data, which shows a slight upward shift in choice probability under the enriched tail configuration and markedly lower probabilities under the long tail configuration. The middle panel presents predictions from EVT-based models, which accurately capture both the shift in the enriched tail configuration and the attenuated responses in the long tail configuration, closely mirroring human behavior. In contrast, the right panel shows that Gaussian models fail to adjust to training distribution changes, producing near-certain predictions on choosing Category B across all configurations. The Gaussian model completely failed for the long tail configuration. This highlights the Gaussian-based models’ inability to model human-like generalization in the presence of skewed category distributions.

Together with results from Experiment 1, these findings reinforce our conclusion that EVT modeling is more effective than Gaussian modeling in capturing changes in human behaviours when it comes to tail-based distributional changes.

#### Statistical tests for normality

To assess whether human responses in the face morph categorization task follow a normal distribution, we applied the Shapiro–Wilk test [Shapiro and Wilk, 1965] to each of the 20 fixed test frames. For each frame and training configuration (uniform, enriched tail, long tail), we gathered 450 binary responses (from 90 morph sequences × 5 participants). Using `scipy.stats.shapiro`, the test was run independently on each group of 90 responses. Across all frames and configurations, p-values were consistently far below 0.05 (e.g., 10^−29^ or lower), indicating strong deviation from normality. These results highlight the limitations of Gaussian assumptions in modeling perceptual decisions and point to the need for alternatives like EVT, which better accommodate asymmetry and individual variability.

To further evaluate the response distributions, we employed the Kolmogorov–Smirnov (K–S) test, which compares the empirical distribution of the data to a normal distribution with matched mean and standard deviation. For each of the 20 test frames, we again used 450 binary responses per configuration and applied the one-sample K–S test using scipy.stats.kstest. All resulting p-values were extremely small (ranging from 10^−50^ to 10^−100^), confirming substantial deviations from normality. These findings align with the Shapiro–Wilk results and further support the use of non-Gaussian modeling frameworks like EVT for capturing the boundary-sensitive, asymmetric patterns observed in human face categorization.

## Conclusion

This work investigates the application of Extreme Value Theory (EVT) as a framework for modeling human visual perception, particularly in recognizing the importance of stimuli near perceptual boundaries. While common models based on Gaussian assumptions emphasize central tendencies in the data, they often overlook the perceptual significance of tail regions — stimuli that can be crucial for recognition and decision-making. We propose that EVT, with its focus on tail distributions, offers a more accurate estimation of human perceptual processes.

To evaluate this hypothesis, we conducted two experiments involving simple line segments and more complex face morph stimuli, each designed to probe human categorization behavior in the gap. Across both tasks, EVT-based models consistently outperformed Gaussian models in capturing the direction of behavioral shifts induced by changes in the underlying stimulus distribution. In particular, EVT was uniquely successful in modeling the effects of tail enrichment and long-tail configurations, aligning more closely with observed human behaviour changes. These findings suggest that EVT provides a more principled and biologically plausible framework for understanding how the visual system prioritizes information near decision boundaries.

## Discussion

One important observation is the divergent effects of the enriched tail configuration across the two experiments likely comes from differences in stimulus structure, task designs, and the interpretive context provided by the distributional manipulation. In the line segment experiment, the enriched tail configuration led to a downward shift in Category B choices, suggesting that participants were less likely to classify stimuli as belonging to Category B when the tail was enriched. A likely explanation lies in the specific structure of the reference stimuli, which consisted of five segments with lengths of [300, 325, 350, 575, 600]. This configuration includes three short segments and only two long ones, resulting in an asymmetric distribution that visually emphasizes the shorter stimuli. Given the simplicity and low-dimensionality of the line segment stimuli, the cluster of short lines likely stood out more saliently and became more strongly associated with Category A during learning. Consequently, when additional rare, long-tail stimuli were introduced during the test phase, they may have been perceived as outliers or as less representative of the more frequent short-length category. This asymmetry could have biased participants toward choosing Category A, thereby reducing the probability of choosing B in the enriched tail configuration.

In the face morph experiment, the enriched tail configuration led to a small upward shift in the likelihood of choosing option B, suggesting that rare stimuli still influenced perception. However, this shift was noticeably smaller than in the line segment task. One possible explanation is that face stimuli are much more complex and high-dimensional, which may reduce the overall impact of adding a few rare examples. In simpler tasks, like the line segments, a few standout examples can strongly shape how people form categories. But in the face task, participants may have relied on a broader sense of the entire reference stimuli set rather than just a few specific faces. It’s also possible that the last face in the reference set (representing Category B) stood out clearly by itself, making the added rare faces less important. Overall, this suggests that enriched tails still affect human perception, but their influence becomes weaker as the task becomes more complex and the category structure more distributed.

These contrasting outcomes demonstrate that enriched tail configurations consistently influence human perception, but the direction of their impact depends on the structure of the task and stimulus space. In some tasks, extreme values may enhance category distinctions, while in others, they may act as perceptual outliers that bias responses away from certain categories. This context-dependent modulation highlights the importance of using models, such as EVT, that are sensitive to distributional extremes and capable of adapting flexibly to different stimulus configurations –— unlike Gaussian models, which assume symmetry and central tendency.

Another observation is in the discrepancy in model performance between the two experiments. This may lie in the nature of the stimuli and the way the reference and testing samples were defined. In Experiment 1, the reference and testing line segments were chosen such that their physical lengths did not overlap, maintaining a clear separation between the two sets. In contrast, in Experiment 2, which involved categorization of morphed human faces, both reference and testing stimuli were selected based on discrete frame indices along a morphing continuum, with no index overlap between the two sets. However, to enable model training and evaluation, these stimuli were subsequently projected onto a compressed one-dimensional Euclidean distance space. After this transformation, reference and testing samples that were originally distinct in terms of frame indices ended up occupying overlapping positions in the distance space. This overlap affected the tail regions of the stimulus distribution, which contains regions that are particularly critical for EVT-based models that rely on the extremes of the distribution to estimate category boundaries. As a result, the degraded distinction between reference and testing sets in the compressed space likely undermined the ability of EVT models to capture the detailed shape of the human categorization curves in Experiment 2, even though they were still able to approximate the overall response trends. Notably, this same compression also impacted the performance of Gaussian-based models, which tended to produce uniformly high Category B probabilities across the testing range in Experiment 2, further indicating that the stimulus representation method may have introduced unintended distortions.

## Limitations and Future Work

One limitation of our work is the reliance on a limited dataset comprising only line segments and human face morphs. While these stimuli are useful for probing perceptual boundaries, they do not fully represent the complexity of real-world visual stimuli, which span a much broader range of features and dimensions. Moreover, the face morph data, in particular, posed additional challenges. To apply our models, we had to compress the high-dimensional face data into one-dimensional points, a simplification that resulted in an unintended aliasing effect. This dimensionality reduction may have affected the accuracy of our modeling results, as the true structure of the data was not fully preserved, leading to suboptimal performance in both EVT and Gaussian models.

As for future work, we believe this work lays the groundwork for integrating EVT into computational models, with potential implications for machine learning algorithms that require strong a generalization property and ability to handle rare events. Future studies would benefit from using more diverse and higher-dimensional datasets to better capture the intricacies of human perceptual processes. An investigation into new EVT algorithms that support multidimensional fitting is also of interest.

## Supporting information

supp_mat

## Acknowledgments

This research was sponsored by the National Science Foundation (NSF) grant CAREER-1942151.

## Footnotes

1 Approved by University of Notre Dame IRB under protocol 18-01-4341. This IRB covers both experiments described in this paper.

2 Code is available at: https://github.com/alyssaq/face_morpher

