## Supplementary material for "Extreme Value Theory for Modeling Category Decision Boundaries in Visual Recognition": supp_mat

### Appendix

#### A Additional Details For Figure 3

Here we include details for the simulation data, which was used to generate the plot in the third row of Figure 3 in the main paper. To generate these illustrative curves, we also used one-dimensional line segments as our input to the models. Table A.1 shows the lengths of line segments used in each setup. The Gaussian Naive Bayes model and the EVT model with probability of inclusion and probability of exclusion were then trained with these data. The test data consisted of discrete morph values ranging from 150 to 300 in steps of 1, representing points along the stimulus continuum used to evaluate model predictions.

| **Configuration** | **Line Segments Used for Category A** | **Line Segments Used for Category B** |
| --- | --- | --- |
| Uniform | 110, 120, 130, 140, 150 | 330, 375, 450, 525, 600 |
| Long Tail | 110, 120, 130, 140, 150 | 300, 400, 500, 600, 700, 710, 720, 730, 740, 750 |
| Enriched Tail | 110, 120, 130, 140, 150 | 300, 375, 450, 600, 800 |

Table A.1: Reference line segment pixel length for category A and category B for the three configurations.

#### B Additional Details For Experiment 1

Here we provide some additional details about our data collection process on human face morph sequences, including screen shots for key steps. The data collection application is very user-friendly. Built with Python and Flask in the back-end and distributed on Amazon Mechanical Turk, the survey can be accessed simply by using a web link.

After a subject accesses the link, they first view an introductory page as shown in Figure A.1. This page includes following information:

1. Description, length and completion criteria. A brief overview about the survey, including the number of questions contained in each survey and what a subject is supposed to do to accomplish it, as well as what happens when a subject finishes answering all the questions.

2. Completion criteria. In order to be approved, the subject needs to submit the survey code given at the end of a survey. A submission will be rejected if any of the following problems are present: (1) Failure to complete the survey. (2) Unable to provide a valid survey code. (3) Multiple submissions of an identical survey code.

3. Troubleshooting guidelines. Information on what to do if an error shows up on the website is provided.

After the subject reads the introduction, the website proceeds to the consent page, as shown in Figure A.2 and Figure A.3, where the subject is provided the information about the study, the purpose of the research, how long it

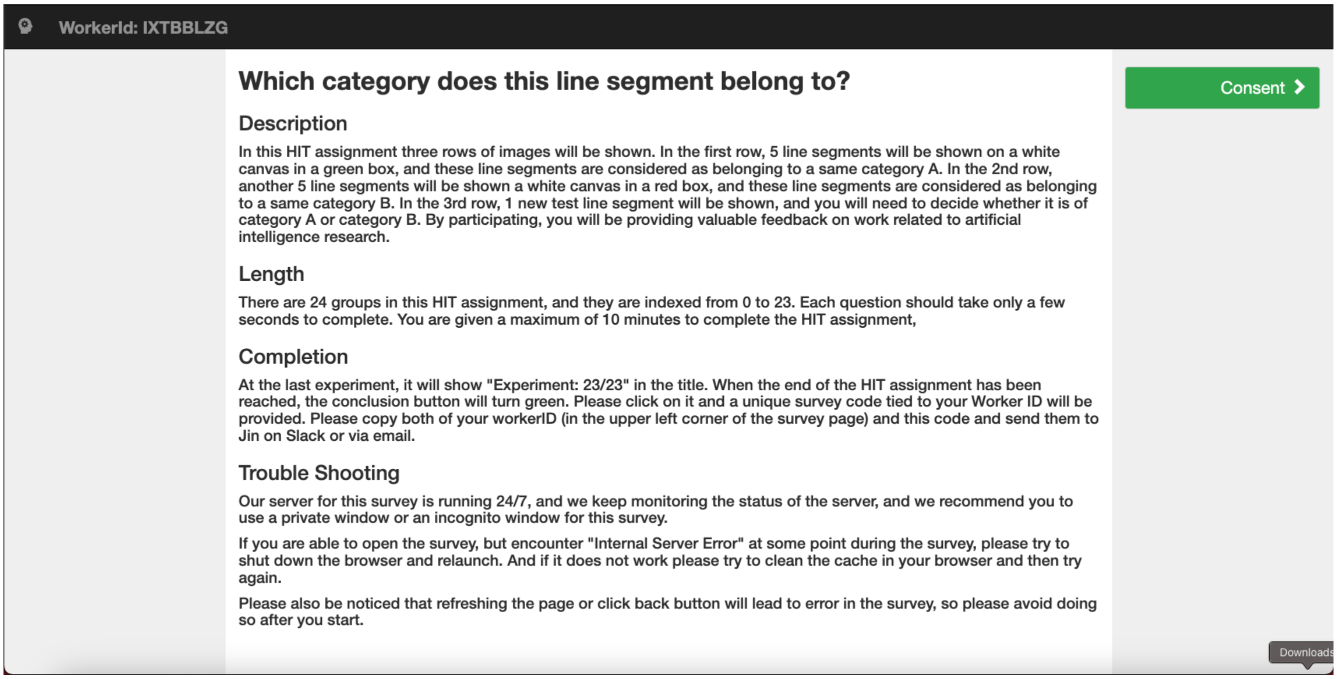


Figure A.1: Introduction page of the survey.

takes to finish a survey, how to end a survey and the privacy policy. The subject must check a box at the bottom of this page to agree to participate.

The survey then moves to the directions page as shown in Figure A.4 and Figure A.5. This page has an overview of the format of the questions that will appear in the survey, and it introduces the question format in detail: (1) showing one image that contains five line segments that belong to category A in the first row; (2) showing one image that contains five line segments that belong to category B in the second row; (3) showing one test segment in the third row and asking a subject to decide whether it is of category A or category B; (4) giving direction on how to make a selection by either clicking on the buttons under an image or pressing a number key on a keyboard.

After the directions, the survey moves to a practice page that has a sample question (as shown in Figure A.6) to make sure that the subject understands how to answer the questions. The survey will only move forward to the real questions after this practice question is answered correctly.

For the actual data collection, 24 survey questions are presented to the subject, who will answer them as each question appears. Figure A.7 shows an example question from the real experiment. After all of the questions are answered, a conclusion page will appear, and a survey code is provided for confirmation of completion.

### C Additional Details For the Data Used in Experiment 2

**Demographic Information.** In our study, we utilized a dataset consisting of 90 morph sequences, strategically distributed across demographic categories to ensure representative coverage. Specifically, the dataset includes 24 morphs for white males, 24 morphs for white females, 29 morphs for Black/Brown males, and 13 morphs for

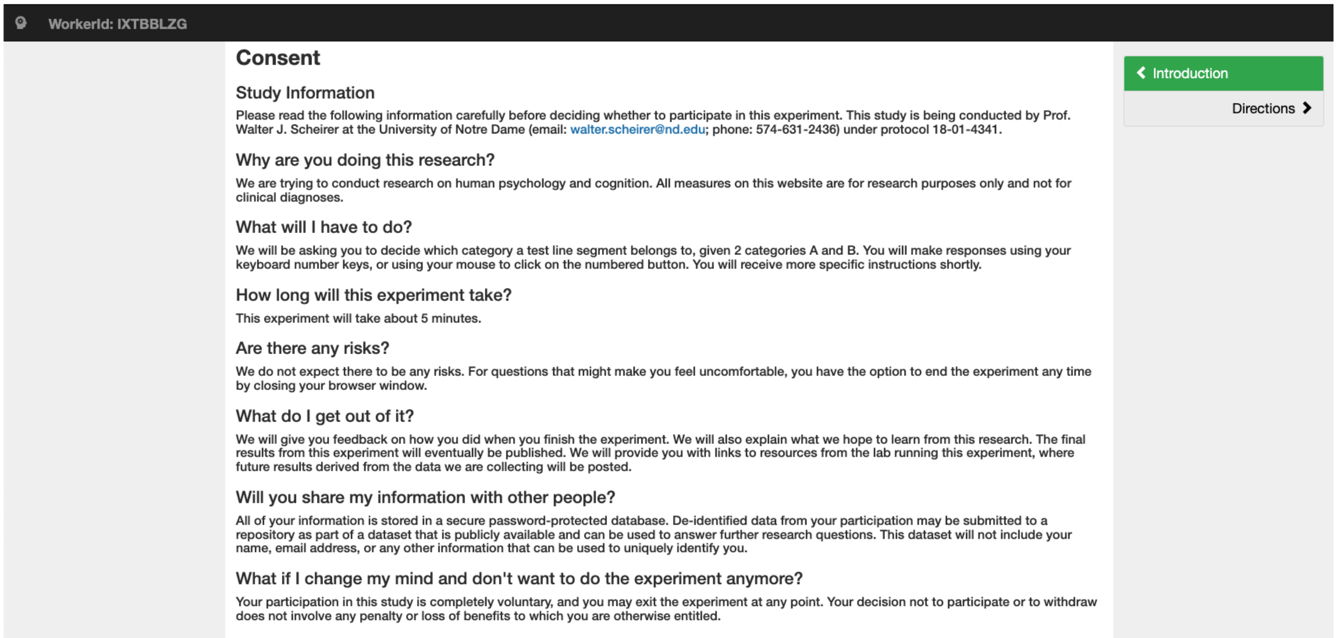


Figure A.2: Consent page of the survey (1/2).

Black/Brown females. This distribution reflects an intentional effort to balance the number of morphs across different gender and ethnicity groups, thereby supporting the robustness and generalizability of our findings. By accounting for these demographic factors, we aim to provide a more comprehensive evaluation of our methods and their applicability to diverse populations.

**Morph Distributions.** Table A.2 presents the distribution of morphs across different skin tones, genders, and views. It includes data for white and black or brown individuals, with male and female categories for both groups. The majority of the morphs are front view images because frontal views typically produce higher-quality and more consistent morphs. Front views offer a clear and unobstructed perspective of the face, minimizing distortions that can occur from angles. This ensures that key facial features, such as the eyes, nose, and mouth, are properly aligned and more easily captured in the morphing process. Additionally, front view images reduce the complexity of factors like head orientation and lighting, which can vary significantly in side or angled views. As a result, the morphing algorithm can more accurately blend facial features when the images are aligned in a front-facing position. This alignment helps preserve the natural look of the morphs and enhances the overall quality, making them more representative of the individuals being modeled.

### D Additional Details For Experiment 2

Similar to experiment 1, we also utilize an application built with Python and Flask in the back-end and distributed on Amazon Mechanical Turk, the survey can be accessed simply by using a web link.

After a subject accesses the link, they first view an introductory page as shown in Figure A.8. This page includes

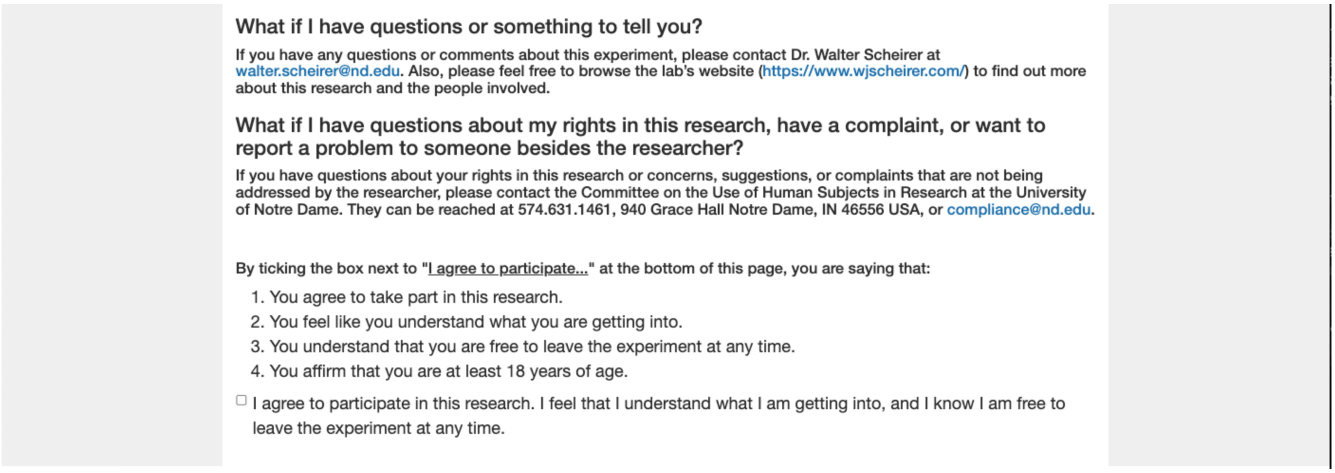


Figure A.3: Consent page of the survey (2/2).

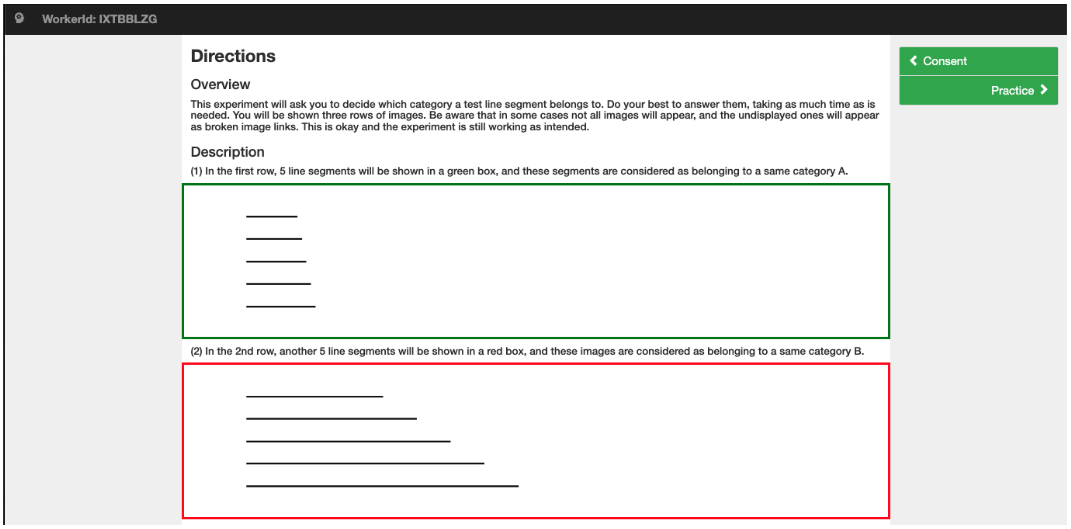


Figure A.4: Direction page of the survey (1/2).

| Skin Tone | Gender | View | Count |
|---|---|---|---|
| White | Male | Front View | 24 |
| White | Female | Front View | 24 |
| Black or Brown | Male | Front View | 22 |
| Black or Brown | Male | Left View | 4 |
| Black or Brown | Male | Right View | 3 |
| Black or Brown | Female | Front View | 9 |
| Black or Brown | Female | Left View | 4 |
| Black or Brown | Female | Right View | 1 |

Table A.2: Distribution of morphs by skin tone, gender, and view.

following information:

1. Description, length and completion criteria. A brief overview about the survey, including the number of questions contained in each survey and what a subject is supposed to do to accomplish it, as well as what

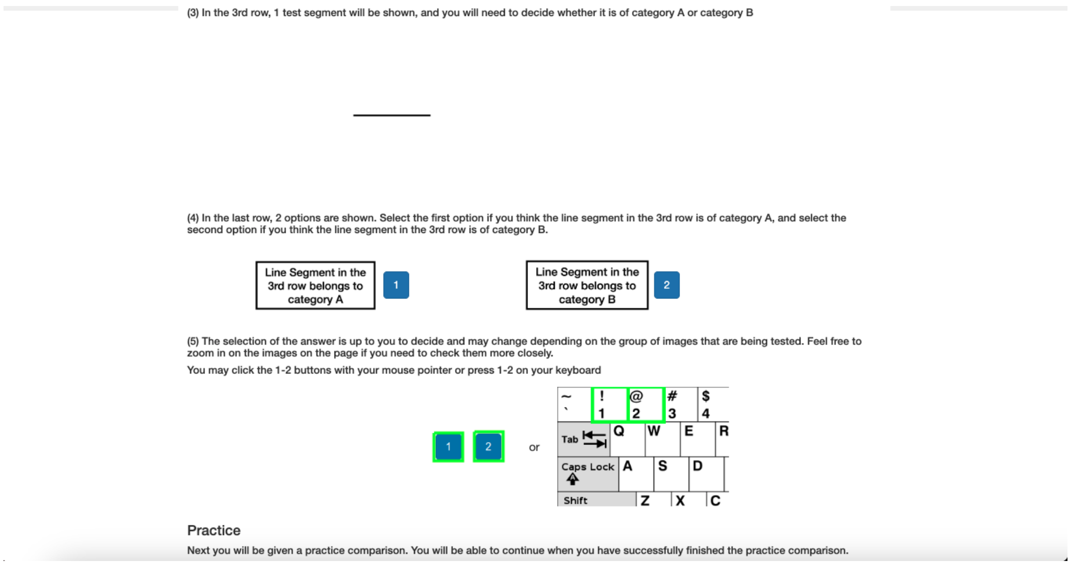


Figure A.5: Direction page of the survey (2/2).

happens when a subject finishes answering all the questions.

2. Completion criteria. In order to be approved, the subject needs to submit the survey code given at the end of a survey. A submission will be rejected if any of the following problems are present: (1) Failure to complete the survey. (2) Unable to provide a valid survey code. (3) Multiple submissions of an identical survey code.

3. Troubleshooting guidelines. Information on what to do if an error shows up on the website is provided.

After the subject reads the introduction, the website proceeds to the consent page, as shown in Figure A.9 and Figure A.10, where the subject is provided the information about the study, the purpose of the research, how long it takes to finish a survey, how to end a survey and the privacy policy. The subject must check a box at the bottom of this page to agree to participate.

The survey then moves to the directions page. This page has an overview of the format of the questions that will appear in the survey, and it introduces the question format in detail: (1) showing 5 images that belongs to person A in the first row; (2) showing 5 images that belongs to person B in the second row; (3) showing one test image in the third row and asking a subject to decide whether it is of person A or person B; (4) giving direction on how to make a selection by either clicking on the buttons under an image or pressing a number key on a keyboard.

After the directions, the survey moves to a practice page that has a sample question to make sure that the subject understands how to answer the questions. The survey will only move forward to the real questions after this practice question is answered correctly.

For the actual data collection, 360 survey questions are presented to the subject, who will answer them as each question appears. After all of the questions are answered, a conclusion page will appear, and a survey code is provided for confirmation of completion.

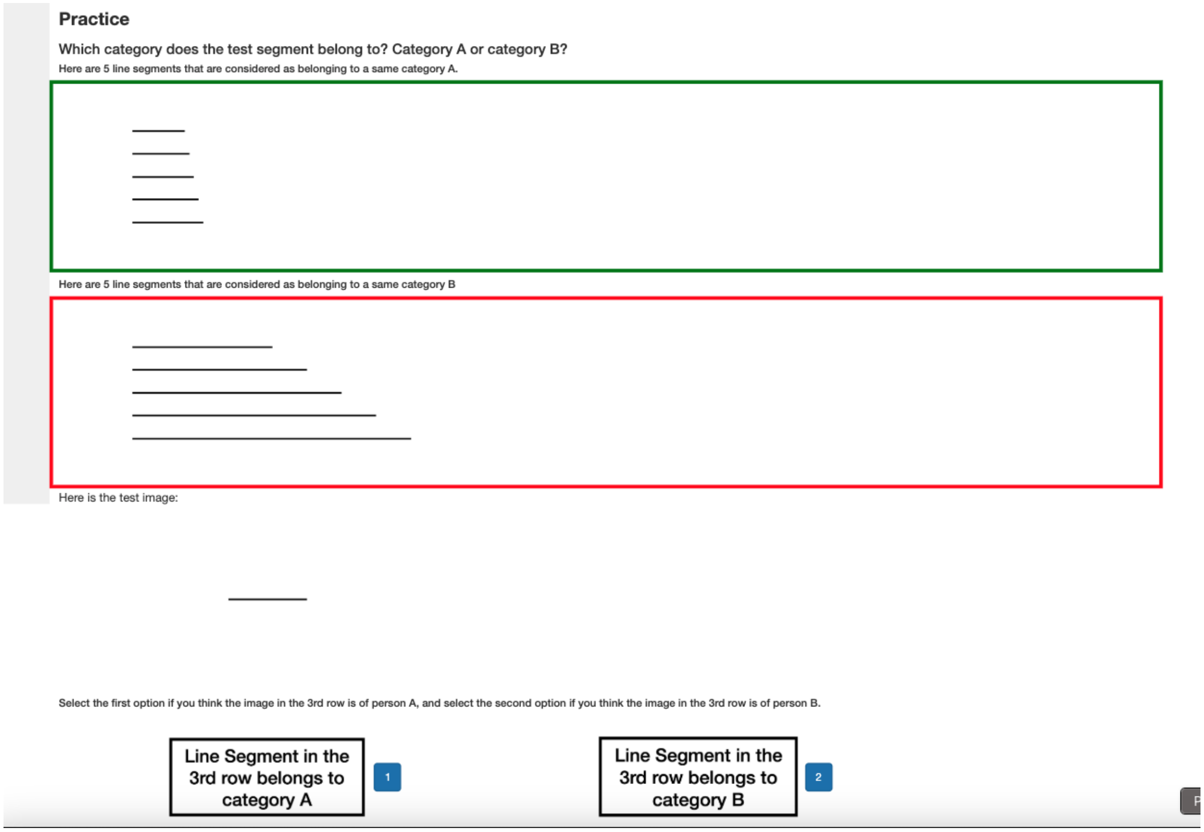


Figure A.6: Practice question.

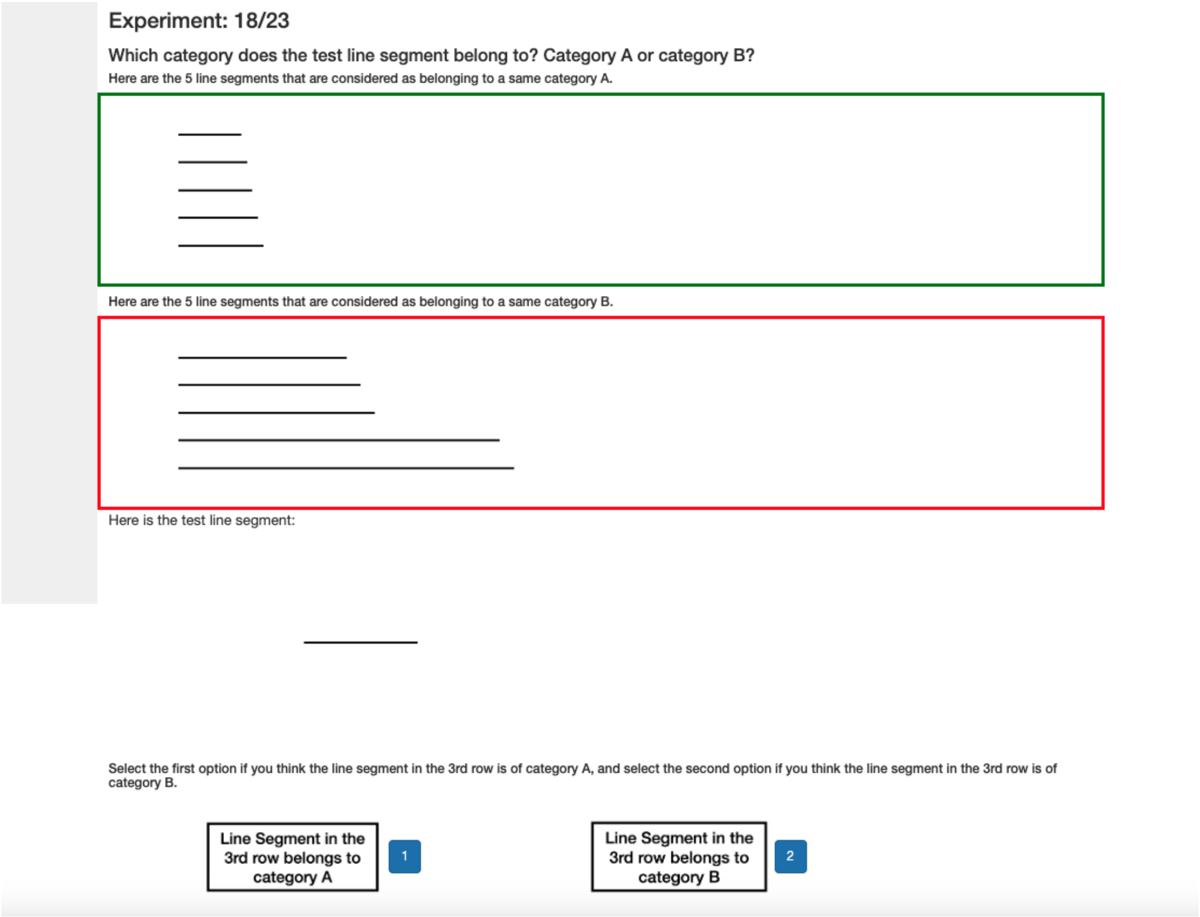


Figure A.7: One example question from the real experiment.

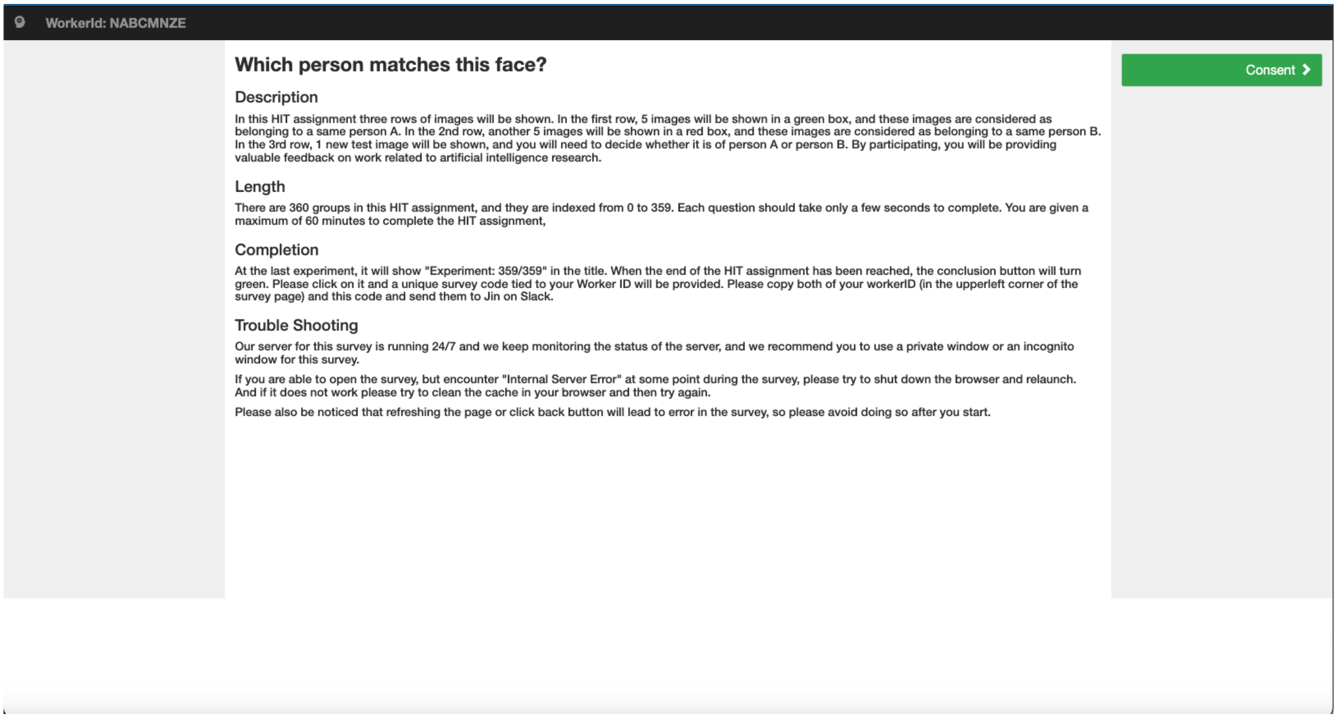


Figure A.8: Introduction page of the survey.

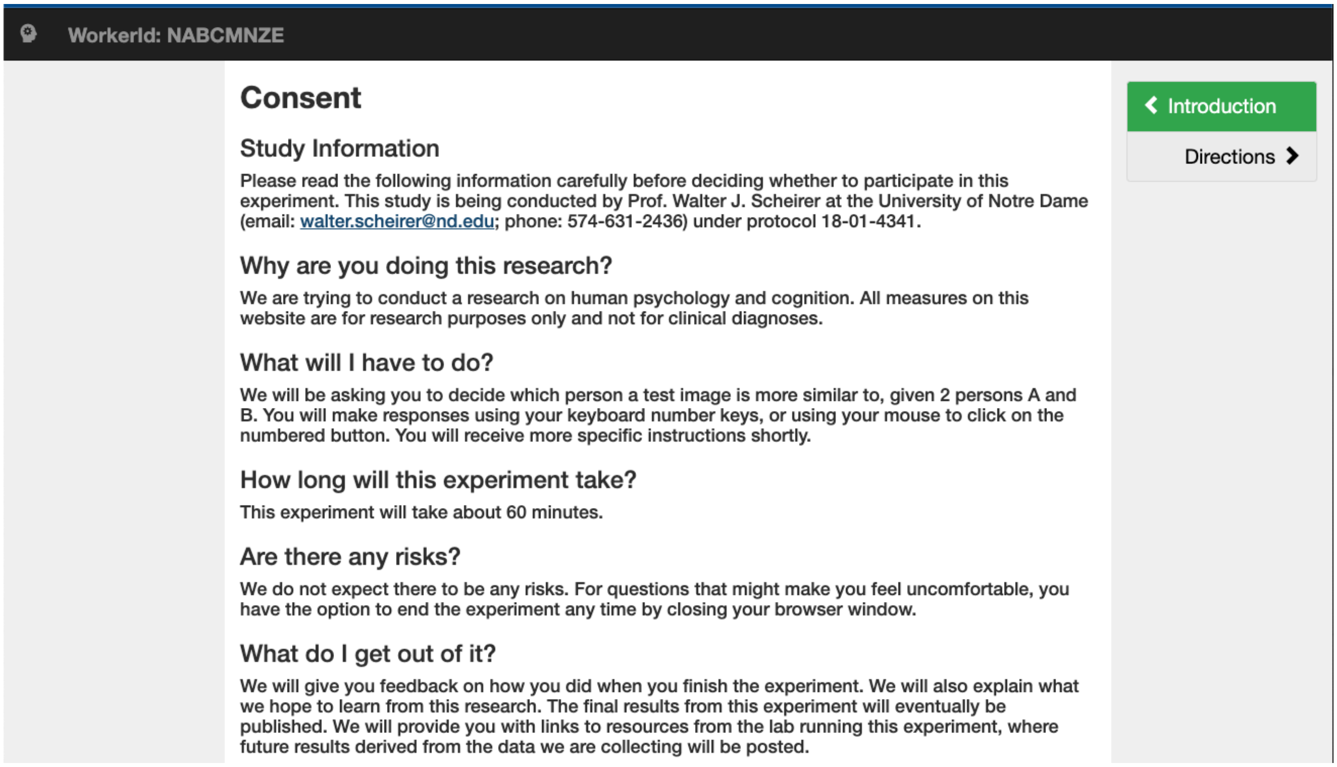

WorkerId: NABCMNZE

### Consent

#### Study Information

Please read the following information carefully before deciding whether to participate in this experiment. This study is being conducted by Prof. Walter J. Scheirer at the University of Notre Dame (; phone: 574-631-2436) under protocol 18-01-4341.

#### Why are you doing this research?

We are trying to conduct a research on human psychology and cognition. All measures on this website are for research purposes only and not for clinical diagnoses.

#### What will I have to do?

We will be asking you to decide which person a test image is more similar to, given 2 persons A and B. You will make responses using your keyboard number keys, or using your mouse to click on the numbered button. You will receive more specific instructions shortly.

#### How long will this experiment take?

This experiment will take about 60 minutes.

#### Are there any risks?

We do not expect there to be any risks. For questions that might make you feel uncomfortable, you have the option to end the experiment any time by closing your browser window.

#### What do I get out of it?

We will give you feedback on how you did when you finish the experiment. We will also explain what we hope to learn from this research. The final results from this experiment will eventually be published. We will provide you with links to resources from the lab running this experiment, where future results derived from the data we are collecting will be posted.

< Introduction

Directions >

Figure A.9: Consent page of the survey (1/2).

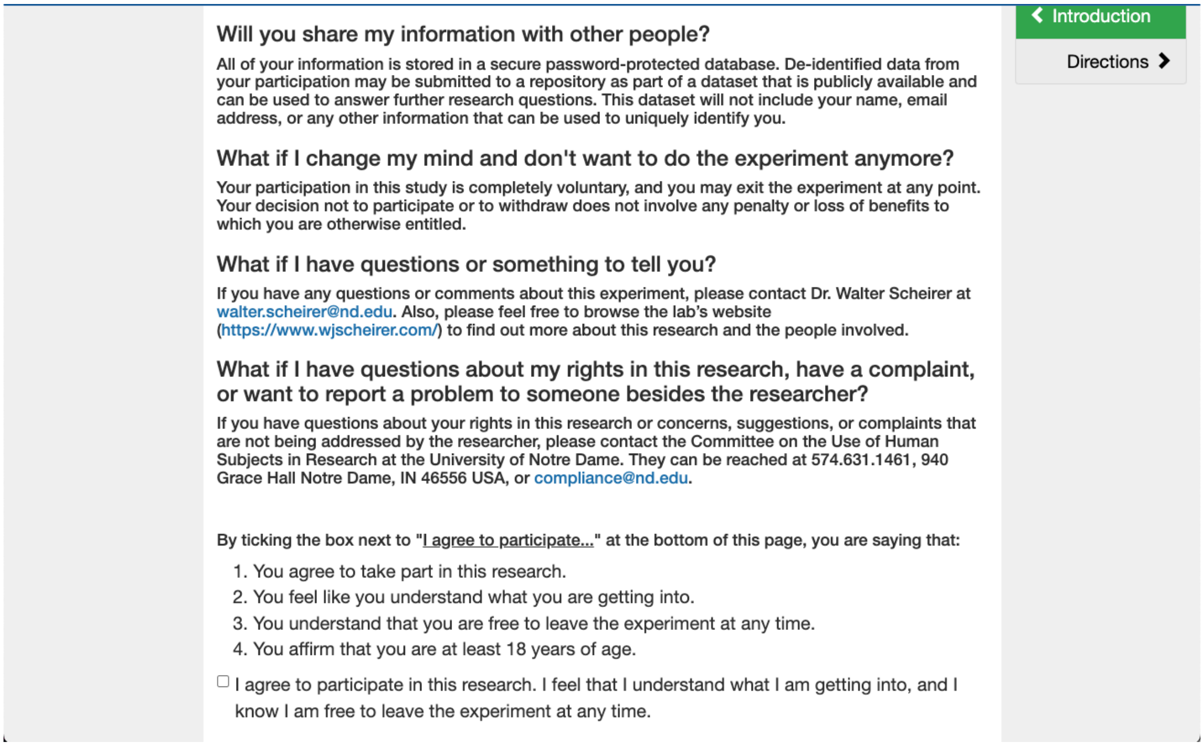

#### Will you share my information with other people?

All of your information is stored in a secure password-protected database. De-identified data from your participation may be submitted to a repository as part of a dataset that is publicly available and can be used to answer further research questions. This dataset will not include your name, email address, or any other information that can be used to uniquely identify you.

#### What if I change my mind and don't want to do the experiment anymore?

Your participation in this study is completely voluntary, and you may exit the experiment at any point. Your decision not to participate or to withdraw does not involve any penalty or loss of benefits to which you are otherwise entitled.

#### What if I have questions or something to tell you?

If you have any questions or comments about this experiment, please contact Dr. Walter Scheirer at. Also, please feel free to browse the lab's website (https://www.wjscheirer.com/) to find out more about this research and the people involved.

#### What if I have questions about my rights in this research, have a complaint, or want to report a problem to someone besides the researcher?

If you have questions about your rights in this research or concerns, suggestions, or complaints that are not being addressed by the researcher, please contact the Committee on the Use of Human Subjects in Research at the University of Notre Dame. They can be reached at 574.631.1461, 940 Grace Hall Notre Dame, IN 46556 USA, or.

By ticking the box next to "I agree to participate..." at the bottom of this page, you are saying that:

1. You agree to take part in this research.
2. You feel like you understand what you are getting into.
3. You understand that you are free to leave the experiment at any time.
4. You affirm that you are at least 18 years of age.

☐ I agree to participate in this research. I feel that I understand what I am getting into, and I know I am free to leave the experiment at any time.

< Introduction

Directions >

Figure A.10: Consent page of the survey (2/2).
